# Human neural dynamics of real-world and imagined navigation

**DOI:** 10.1101/2024.05.23.595237

**Authors:** Martin Seeber, Matthias Stangl, Mauricio Vallejo, Uros Topalovic, Sonja Hiller, Casey H. Halpern, Jean-Philippe Langevin, Vikram R. Rao, Itzhak Fried, Dawn Eliashiv, Nanthia Suthana

## Abstract

The ability to form episodic memories and later imagine them is integral to the human experience, influencing our recollection of the past and our ability to envision the future. While research on spatial navigation in rodents suggests the involvement of the medial temporal lobe (MTL), especially the hippocampus, in these cognitive functions, it is uncertain if these insights apply to the human MTL, especially regarding imagination and the reliving of events. Importantly, by involving human participants, imaginations can be explicitly instructed and their mental experiences verbally reported. In this study, we investigated the role of hippocampal theta oscillations in both real-world and imagined navigation, leveraging motion capture and intracranial electroencephalographic recordings from individuals with chronically implanted MTL electrodes who could move freely. Our results revealed intermittent theta dynamics, particularly within the hippocampus, which encoded spatial geometry and partitioned navigational routes into linear segments during real-world navigation. During imagined navigation, theta dynamics exhibited similar, repetitive patterns despite the absence of external environmental cues. Furthermore, a computational model, generalizing from real-world to imagined navigation, successfully reconstructed participants’ imagined positions using neural data. These findings offer unique insights into the neural mechanisms underlying human navigation and imagination, with implications for understanding episodic memory formation and retrieval in real-world settings.

## Introduction

Human cognition is a complex interplay of processes, with the formation of episodic memories and the imaginative capacity to revisit, manipulate, and extrapolate from these memories representing two of its most fundamental aspects. These cognitive functions lie at the core of human experience, shaping our recollection of past experiences and our ability to construct visions of the future. While a significant portion of research focuses on understanding the neural dynamics underlying these processes in the context of spatial navigation in freely moving rodents, it is widely held that the medial temporal lobe (MTL), particularly the hippocampus, plays a central role ^1,2^. Previous research in rodents has identified “place cells” tuned to specific locations, forming neuronal sequences spanning entire movement^3–6^. Importantly, these neuronal sequences persist during immobile periods^7–10^, implying internal generation and making them well-suited for organizing episodic memories^3,11,12^, mentally simulating these memories ^13^, and planning future behaviors^3,14^. However, it remains unclear whether such mechanisms exist in humans during real-world ambulatory navigation and especially during overt, on-demand imagination or re-experiencing of episodic memories.

To coordinate hippocampal neuronal sequences^15–19^, theta oscillations within the frequency range of ∼4 to 12 Hz^20^ hold significance, as their disruption can lead to spatial memory impairments^2,21,22^. In humans, hippocampal theta oscillations manifest in brief bouts rather than continuously, as observed in rodents, during both virtual and real-world navigation^23–26^. This cross-species difference raises uncertainty regarding whether these intermittent bouts in humans can adequately support neuronal sequences and subsequent segmentation of distinct episodic events during real-world navigation. Furthermore, whether hippocampal theta bouts can be internally generated and segment episodic memories, such as during imagination, remains entirely unknown. Identifying such mechanisms would provide first-in-human insights into the shared neural organization principles between real-world spatial navigation and episodic memory^3,11^, effectively bridging decades of findings across species and integrating spatial navigation and episodic memory research.

In the current study, we compared theta dynamics in the human MTL between real-world and imagined navigation. Imagined navigation involved episodic memories characterized by distinct temporal features, facilitating a direct comparison to real-world navigation. To explore this, we examined intracranially recorded neural oscillations from five participants who had undergone chronic implantation of the responsive neurostimulation device, RNS System (NeuroPace, Inc.), for the treatment of epilepsy^27^. Specifically, we directly compared MTL theta dynamics during real-world navigation to periods where participants mentally simulated navigating the exact same routes while walking on a treadmill.

We observed theta dynamics within the MTL that varied depending on the participant’s position within segments of the navigational routes, effectively encoding the spatial geometry of those routes. These theta dynamics demonstrated temporal consistency across individual trials, were consistently observed in all participants, and occurred during imagined navigation while walking on a treadmill. We further demonstrated the feasibility of estimating relative positions within navigational route segments using theta dynamics of real-world navigation and applied this model to accurately reconstruct imagined positions.

## Results

### Theta dynamics during real-world navigation

We investigated whether neural dynamics in the human MTL are influenced by position, time, or progress on specific spatial routes, building upon previous research^3,28,29,7,30,6,11,9^. To achieve this, we tasked participants with walking along two distinct spatial trajectories (Fig. 1 A-D). Since the angles and locations of turns on these routes were not visible during real-world navigation, participants had to learn and remember the walking patterns, with turns serving as critical retrieval points. Theta dynamics (3-12 Hz) were extracted from iEEG data recorded from chronically implanted electrodes (Fig. 1 C, D). All participants were able to successfully learn the navigational routes as instructed (Fig. 2A, Fig. S1). By synchronizing and aligning iEEG and motion capture data, we were able to compare theta dynamics with the positions along each spatial route.

**Fig. 1.**
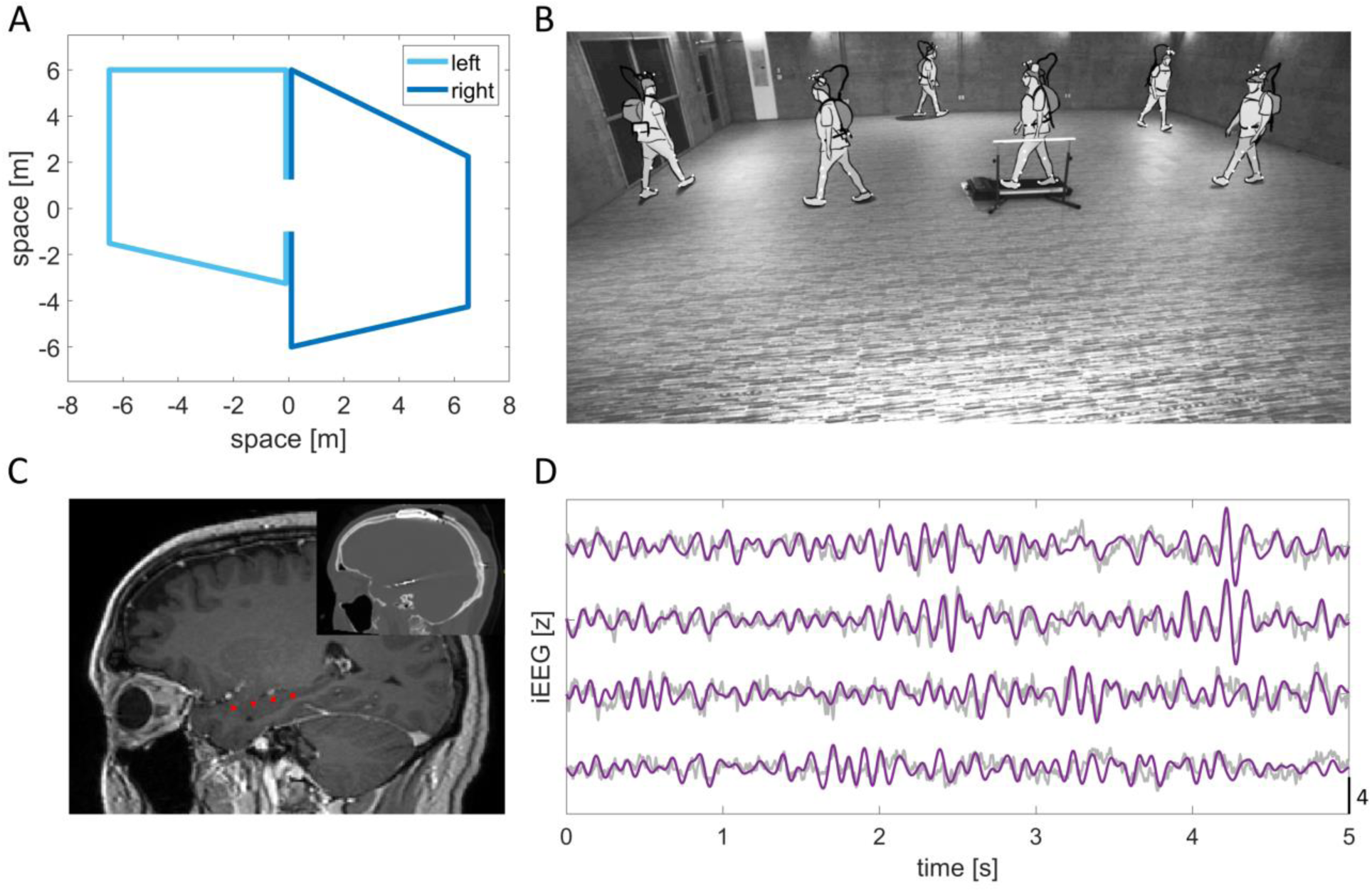
| Experimental paradigm. (**A**), Participants completed a spatial navigation task that involved the learning of two distinct routes: a leftward route represented in light blue and a rightward route represented in dark blue. (**B**), An illustrative time-lapse motion capture shows an exemplified participant navigating the rightward route. Treadmill walking trials were interspersed with real-world walks. During treadmill walking, participants were instructed to mentally simulate their previous or upcoming route. Initially, before their real-world navigation walks, participants engaged in treadmill walking without the additional component of imagined navigation. (**C**), Pre-operative magnetic resonance image and post-operative computed tomography (inset) were used to identify the location of intracranial electrode contacts. Electrode locations of four electrode contacts (comprising two bipolar channels) in the hippocampus from an example participant are indicated in red. (**D**), Intracranial EEG (iEEG) was continuously recorded during both real-world and imagined navigation. Exemplary broadband iEEG activity (gray) during real-world navigation is superimposed with the filtered signal (magenta) within the theta band (4-12 Hz).

**Fig. 2.**
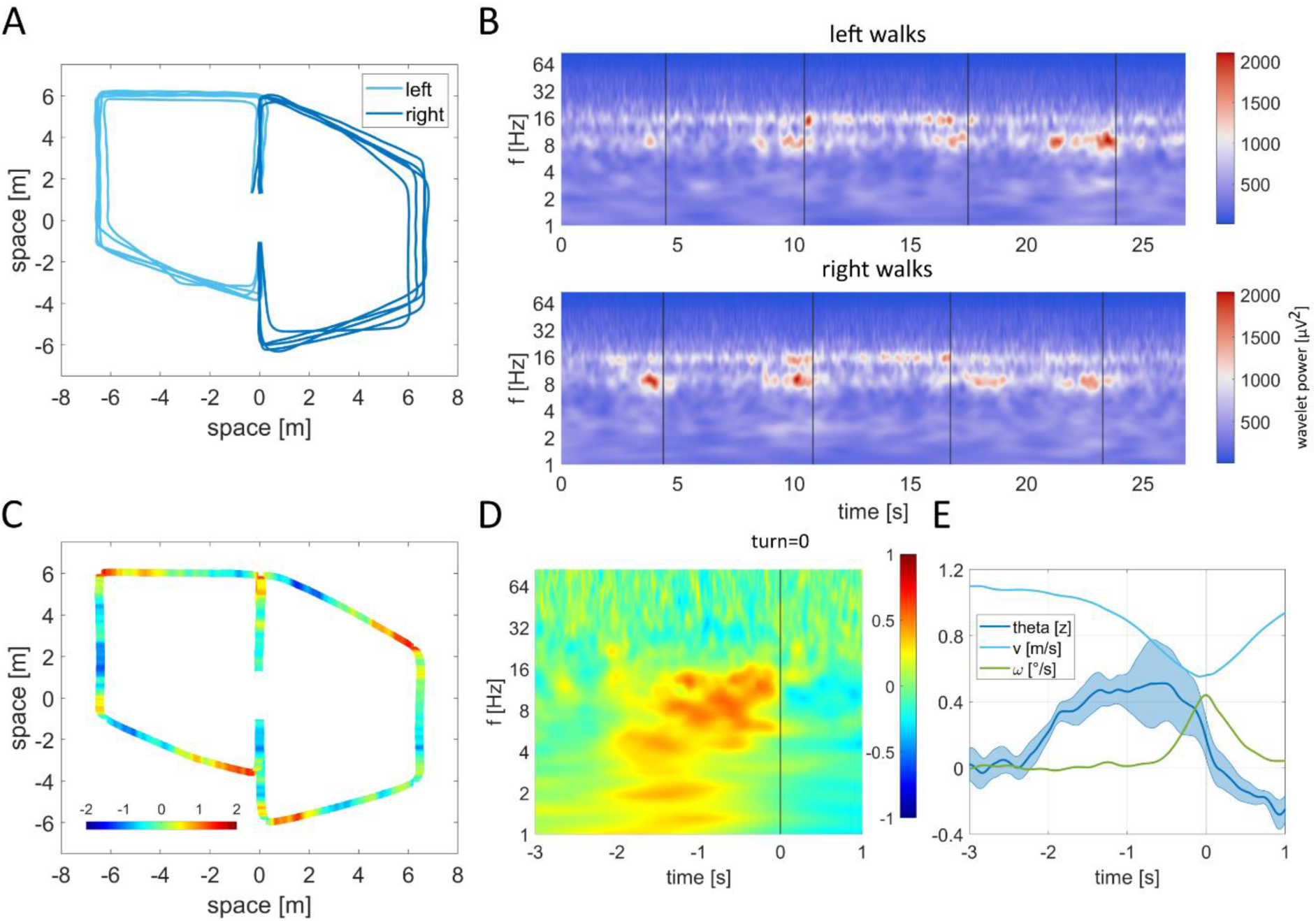
| Theta dynamics during real-world navigation. (**A**), Distinct lines illustrate the average walking routes of each individual participant. (**B**), Time-frequency plots of exemplary hippocampal activity reveal task-related and temporally organized theta oscillations during both left and right walks (top and bottom panels, respectively). Black vertical lines demarcate where turns occured in the routes. (**C**), Theta activity (z-scored) within the left anterior hippocampus (group average, one channel per participant) is overlaid onto the motion trajectories, demonstrating heightened theta power as participants approached upcoming turns. (**D**), Average time-frequency activity aligned with all turns confirms the engagement and the temporal relationship of theta activity preceding turns (time = 0). (**E**), Mean theta activity ± standard error (for frequency ranges see Table 1) juxtaposed with the speed and angular velocity of the hips, averaged across the group and aligned to turns (time = 0).

We observed an increase in MTL theta amplitudes at upcoming turns just before the completion of a linear segment within a walking route (Fig. 2B-D). This pattern remained consistent across all five participants while they navigated both left and right walking routes (Fig. S3, 6A). Interestingly, participant 2 consistently incorporated an additional turn not originally part of the route, and this effect was still observed. We further confirmed that theta activity aligned to route segments through time-frequency analyses (Fig. 2D). For theta frequency ranges at the subject-specific level see Table 1 and methods for details. Notably, theta activity reached its peak significantly before the actual physical turns occurred (–0.93 ± 0.77 s, p < 0.001), as captured by hip rotation (Figure 2E, Fig. S4C). As participants approached a turn, they naturally reduced their walking speed. Concurrently, there was a significant increase in theta power preceding this deceleration, and this pattern was consistent with other behavioral variables, including head rotation, body turning, and eye movements (Fig. S8, S9, Table S2). Thus, these theta dynamics cannot be easily attributed to confounding factors such as walking speed, heading, hip rotation, or eye movements.

Consistent with previous findings^23–26^, theta activities appeared in brief bouts rather than as continuous oscillations, as seen in rodents. We analyzed the timing of these bouts on a single-trial level and observed their alignment with upcoming turns (Fig. 3B). Strikingly, the percentage of trials in which theta bouts occurred at specific time points closely mirrored the trial-averaged theta dynamics (Fig. S2). This finding suggests that theta bouts exhibited task-related alignment across trials. More specifically, these bouts aligned with particular instances along the navigational routes, effectively dividing participants’ movement trajectories into distinct linear segments.

**Fig. 3.**
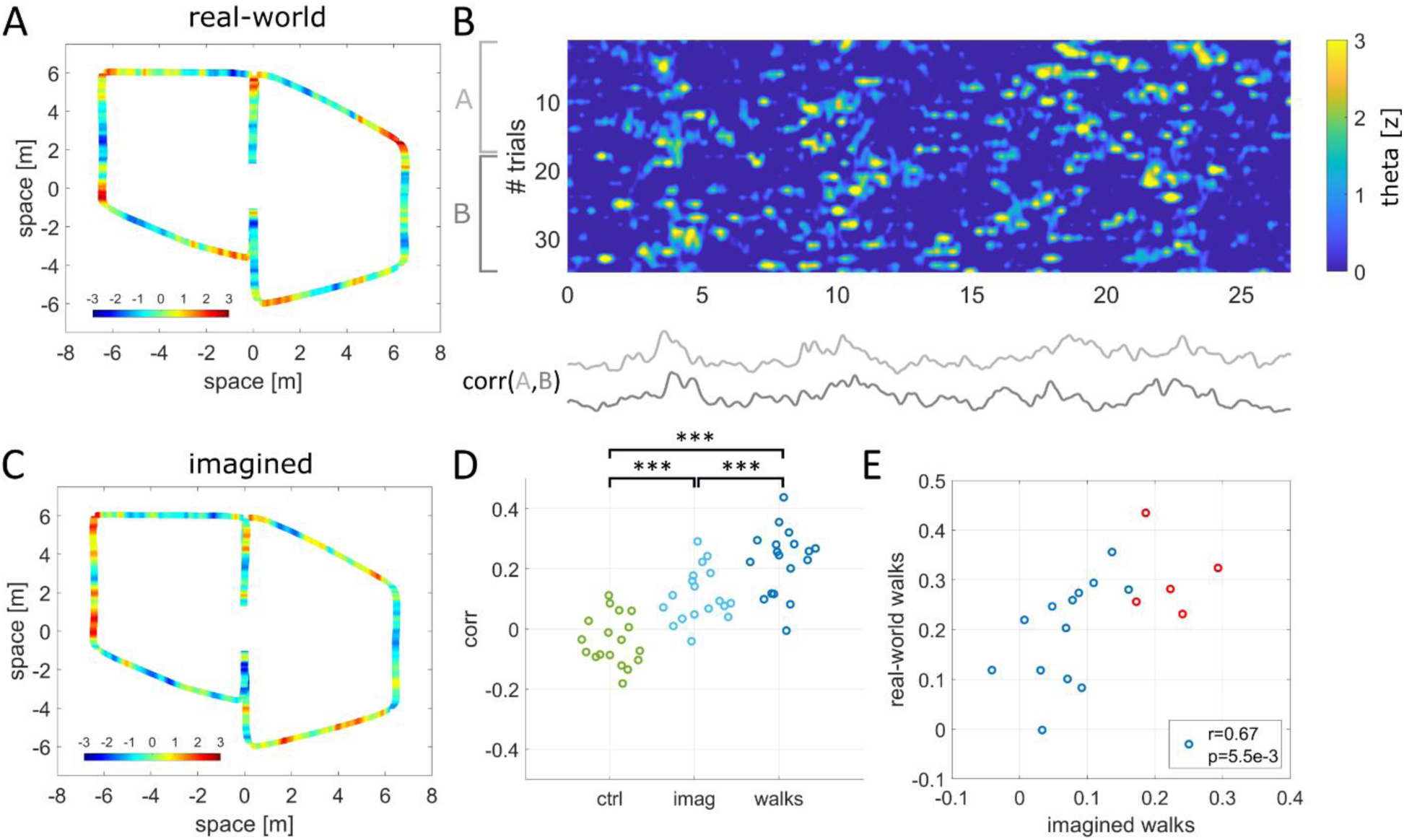
| Comparative theta dynamics of real-world and imagined navigation. (**A**), Theta activity (weighted sum of all channels) averaged across the group during real-world navigation superimposed on the mean motion trajectory of all participants. (**B**), The temporal consistency of theta dynamics was assessed by calculating the correlation between mean signals obtained from randomly dividing the data (trials) into two halves (A and B). (**C),** Theta activity (weighted sum of all channels) averaged across the group during imagined navigation rendered on the group mean motion trajectory. Note that theta activity tended to increase earlier during imagined turns compared to real-world turns. (**D**), Temporal consistency of theta dynamics was significantly higher during both imagined and real-world navigation when compared to sole treadmill walking. Data from each of the 18 intracranial electrode channels is represented by individual circles. (**E**), Notably, temporal consistency was spatially correlated between real-world and imagined navigation. Electrodes showing higher consistency in theta dynamics during real-world navigation also demonstrated comparable consistency during imagined navigation, suggesting the presence of analogous functional networks engaged in both types of navigation. Interestingly, the highest consistency electrodes were located in the left anterior hippocampus (red circles), in contrast to other regions within the MTL (blue circles).

### Shared neural dynamics in real and imagined navigation

The dynamics within the hippocampus are known for their capacity to encode task-related structures associated with position, time, and, in a broader sense, sequences. Building upon the knowledge that hippocampal neuronal dynamics can be internally generated^7,9,8,10^, we investigated whether MTL oscillatory dynamics exhibit similarities during real-world and imagined navigation. To explore this, we incorporated intervals of real-world navigation trials interspersed with treadmill walking periods. During the initial treadmill walks (control trials), participants were not given any explicit instructions other than to walk at a steady speed. However, on the later treadmill walks (imagination trials), they were instructed to recollect their previous or upcoming routes while walking at the same steady speed. Each treadmill condition consisted of 24 trials, resulting in a total of 72 trials (24 trials each for control, imagination left route, imagination right route, Table S1).

Given the pronounced presence of task-related structured theta dynamics during real-world navigation, we explored whether analogous temporal patterns were evident during imagined navigation. Our analysis showed that the temporal consistency across trials (Fig. 3B, Fig. S6B) was significantly higher for theta dynamics during imagined navigation (p < 0.001) compared to the control condition of sole treadmill walking (Fig. 3D). The highest level of temporal consistency was observed during real-world navigation, whereas this consistency was absent during sole treadmill walking (p = 0.96). We did not find an effect of the time reference (previous/upcoming route) of imaginations but a modest (p = 0.023) session effect, consistent with learning throughout the experiment (Fig. S5A, B). Next, we examined the congruence of functional anatomy within the MTL during both real-world and imagined navigation. Our analysis revealed a spatial correlation in the temporal consistency values (r = 0.67, p = 0.006) between these conditions. Notably, recording channels demonstrating heightened temporal consistency during real-world navigation, such as within the left anterior hippocampus, were found to manifest a similar pattern during imagined navigation, suggesting the involvement of comparable functional networks in both modes of navigation (Fig. 3E).

As a means to validate participants’ engagement in imagining the distinct navigational routes, we examined their eye movements. During real-world navigation, participants tended to focus their gaze ahead on the walking path, resulting in relatively higher probabilities of left or right gazes for leftward and rightward walks, respectively. While this effect was less pronounced during imagined navigations, subtle eye movements still differentiated between imagined left and right routes (Fig. S7). However, we did not find a direct effect of eye movements on theta amplitudes during imaginations (p = 0.58). Instead, we observed that theta dynamics were primarily related to the relative position within the route segments (p = 0.006), mirroring the patterns observed during real-world navigation. The impact of relative position during imagined navigation was significantly larger than that of eye movements (p = 0.003), indicating that eye movements did not drive the effect (Fig. S9C, D). We next explored the feasibility of estimating positions of real-world walks from theta dynamics and assessed whether such reconstructions could generalize to imagined navigation.

### Reconstructing imagined positions

We examined the feasibility of reconstructing participants’ route progression based on neural activity patterns. This exploration was motivated by the observation that both real-world and imagined navigation induced temporally structured theta dynamics within the MTL. To this end, we investigated whether these theta dynamics encoded the inherent task structure of the routes as illustrated in Fig. 4A. These routes were designed with five linear segments connected by four turns. Upon accounting for variations in segment lengths, it became evident that theta amplitudes were consistently modulated across the segments. However, distinct recording electrodes exhibited peak activations at slightly varying positions within a route segment (Fig. 4C). In light of this finding, we developed a model that considered the unique timings of each electrode to capture the relative positions of route segments, treating them as two-dimensional circular variables, a methodology based on prior research^31^. Utilizing a linear regression model, we used theta dynamics from all MTL channels as predictors and using the task structure (comprising cosine and –sine phase alignment) as response variables. Regression coefficients were learned from subsets of the data, and the model’s ability to generalize to unseen trials was tested (using 10 x 10 cross-validation) to prevent overfitting. Our results indicated that relative segment positions could be reconstructed better than chance (p < 0.001) during real-world navigation. The performance of the reconstruction was evaluated as normalized probability densities for each relative position within a segment. The time-resolved reconstruction revealed that position estimates matched not only at upcoming turns but encompassed the entire segment of a route (Fig. 4D). Overall reconstruction errors were quantified as the angle between actual and estimated positions because relative positions were treated as circular variables (Fig. 4D).

**Fig. 4.**
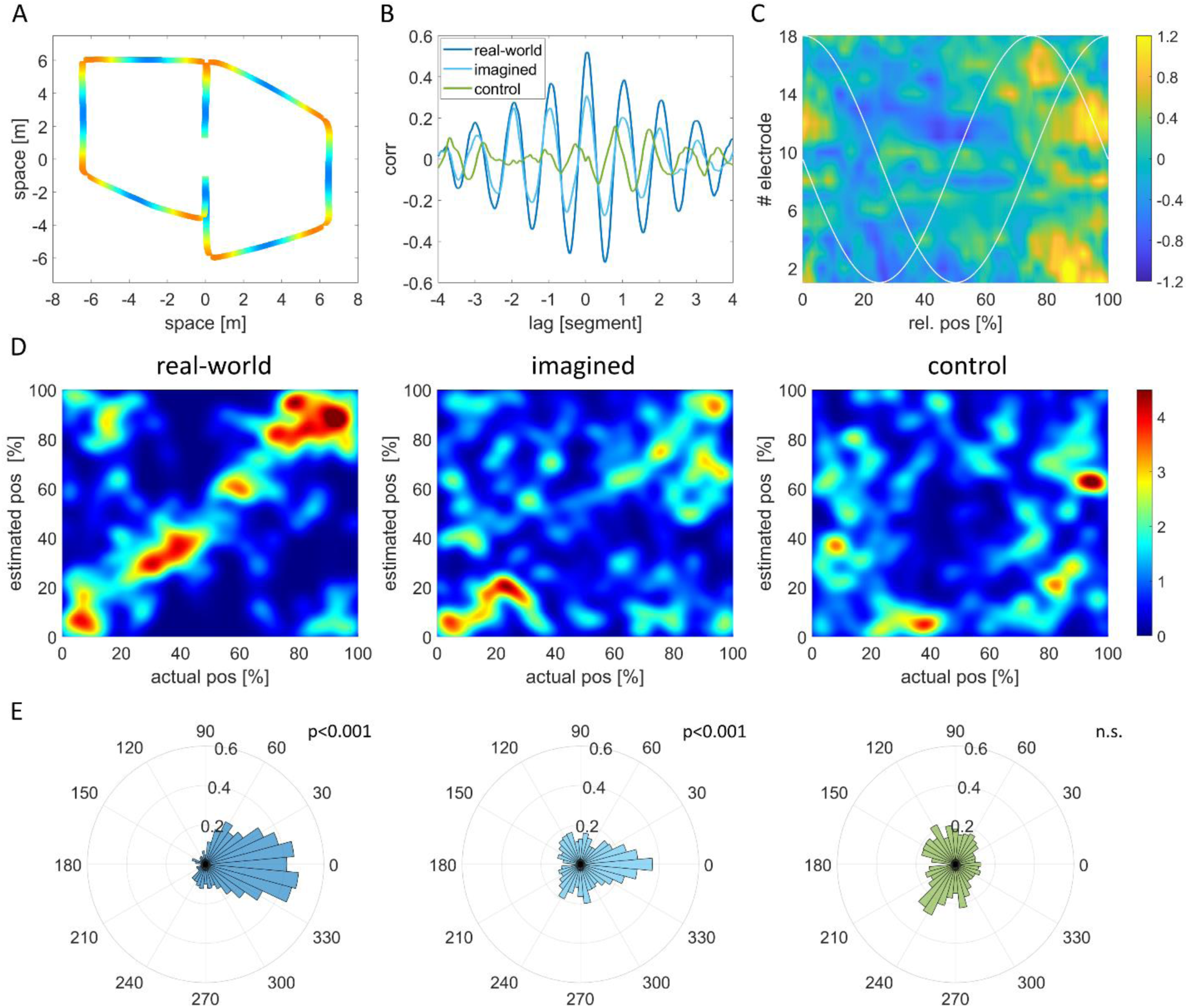
| Reconstructing relative position from neural dynamics. (**A**), Model of the navigational maze structure represented as a sinusoidal pattern peaking at turns. (**B**), Cross-correlation analysis between theta dynamics and the maze structure revealed a significant correlation (lag = 0) for both real-world and imagined navigation, while sole treadmill walking did not exhibit such a correlation. The similarity and repetition of theta dynamics in each maze segment led to four lateral peaks in the cross-correlations at lags that matched the length of one segment on a walking route. (**C**), Theta dynamics are illustrated as a function of relative position within all segments across all 18 electrode channels. Notably, the timing of theta dynamics varied across channels, roughly following a cosine or sine wave pattern (white lines). These two orthogonal theta modulations effectively encoded relative position as a circular variable. (**D**), Reconstruction of relative segment positions from theta dynamics depicted using 2D histograms that show both the actual and estimated positions. Color-coded representations illustrate the probabilities of all possible combinations of actual and estimated positions. An ideal reconstruction outcome would manifest as a diagonal pattern. During real-world navigation, the estimated positions (cross-validated) clustered closely around the actual physical positions within each segment (left panel). During imagined navigation, the estimated positions also aligned consistently with the positions estimated from the duration of the imagination periods particularly at the beginning and end of each route segment (mid-panel), illustrating heightened accuracy in those instances. However, during sole treadmill walking where imagined navigation was absent, reconstruction failed to yield accurate results (right panel). (**E**), Histograms depicting the errors in reconstruction (measured in degrees as a circular variable) for each condition revealed distinct patterns. In both real-world (left panel) and imagined (middle panel) navigation conditions, the errors clustered around zero, indicating accurate reconstruction. However, during sole treadmill walking (right panel), this clustering around zero was absent, signifying less accurate reconstruction.

We applied the position reconstruction model, which was initially trained on real-world walking data, to imagined navigation data. In real-world walking, position and time are linearly related due to well-defined walking velocities. However, in imagined navigation, the imagined velocity is not directly available. We also anticipated that the timing of imagination may not precisely match that of real walks. While the task structure remains consistent during imagination, route segments may undergo stretching and bending. To address this, we aligned estimated imagined positions and the task structure using linear time warping. After alignment, we found that the position estimation model effectively generalized to imagination trials not used in the model’s training and alignment procedure (10 x 10 cross-validation). The reconstruction errors for imagined positions were significantly smaller (p < 0.001) than those derived from the control condition, where imaginations were absent but physical behavior was identical (Fig. 4 D, E). Theta dynamics exhibited a significant correlation between real-world and imagined navigation (r = 0.30, p = 0.003) and were related to the maze structure (Fig. 4A) for both real-world (r = 0.51, p < 0.001) and imagined navigation (r=0.29, p = 0.010), but no such correlation was observed during sole treadmill walking (r = –0.01, p = 0.528). Cross-correlations between theta and maze geometry revealed four lateral peaks at multiple integers of the segments’ lengths, reflecting the four turns on each walking route (Fig. 4B). These results indicate that theta dynamics within distinct segments were similar and repeated themselves along the entire movement trajectory when accounting for varying segment lengths.

## Discussion

Using rare mobile iEEG recordings in freely-moving human participants, we demonstrated that intermittent theta dynamics in the human MTL are well suited to internally organizing the recollection of past episodes or imagination of future behaviors. Specifically, our findings revealed that transient theta oscillations aligned at upcoming turns during real-world navigation, moments which served as critical retrieval points essential for transitioning between navigational route segments. These dynamics exhibited a similar temporal pattern during episodic imagination of previous and upcoming routes. The presence of comparable characteristics during both real-world and imagined navigation suggests neural progression along abstract geometries delineating multipartite imagined routes into more elementary sections. Notably, this repetitive temporal structure was absent when individuals engaged in treadmill walking alone, where mental recapitulation of previously learned spatial routes was absent despite identical physical behavior.

Our finding of structured theta bouts during imagined navigation suggests that hippocampal networks can internally generate neural dynamics representing distinct sections of a task. This task structure may involve sequences of route segments linked by action points, such as turns observed in real-world navigation or event timing during imagination and cognitive tasks with non-spatial components. These findings align with prior rodent studies demonstrating the ability of hippocampal networks to internally generate neuronal sequences^7,8^, with theta oscillations crucial for their temporal organization^15–19^. It is conceivable that the transient theta oscillations we report here in humans could reflect neuronal populations overrepresenting forthcoming turns, marking the transitions between segments. Indeed, hippocampal place cells and other spatially-tuned cell types can bias their activity toward behaviorally relevant locations or moments^32,9,10^. In navigational tasks featuring repetitive segments, such as ours, place cells and grid cells tend to form repetitive firing sequences that reset at turning points within a maze^30^. Task-related theta dynamics observed during imagined navigation also complement recent research suggesting that hippocampal theta oscillations play a key role in dynamically exploring locations related to both current and hypothetical behaviors in rodents^18,33^. Internally generated dynamics such as these are particularly relevant for episodic memory retrieval, where sensory input may be minimal.

These findings also resonate with studies on event boundaries delineating ongoing experiences^34–36^. In our current study, turns can be interpreted as event boundaries, which are predictable since they are defined by participants’ actions. This partitioning of navigational routes into segments also deepens our understanding of the human MTL, showing theta oscillations can adhere to abstract maze geometry rather than physical environmental constraints. Such partitioning might enable flexible combination of previously experienced fragmented episodes to adapt to dynamic real-world task demands. Moreover, the reassembly of neural sequences through transitions might facilitate hippocampal circuitry to generate novel trajectories relevant to envisioning future behaviors. Theta bouts preceding turns also complement prior findings of theta bouts occurring after cognitive boundaries during movie watching^37^. While self-generated actions are predictable, the cuts between movie scenes are not, explaining the temporal alignment of theta bouts before versus after such boundaries in proactive versus passive behavioral paradigms.

Recent studies on simulated navigation have indicated that memory is a primary driver of hippocampal theta oscillations in humans^38^. Our study goes further to describe the fine-grained temporal structure of theta dynamics, which partitioned participants’ trajectories into discrete segments during real-world and imagined navigation. We did not find differences in the temporal consistency of imagined previous versus upcoming routes. These results suggest that MTL dynamics are similar between actual, recapitulated, and possible future behaviors. It is worth noting that we instructed participants to imagine their movements along the navigational routes on the same time scale as during real-world navigation. Episodic memory recall is typically temporally compressed^38^, potentially leading to more events and theta bouts per second, resulting in elevated average theta power. Moreover, increased theta power has also been reported during spatial memory retrieval in stationary, view-based navigational tasks^39,40^. Consistent with functional neuroimaging research on egocentric navigation strategies ^41^, sequence representation^42^, and imagination of events^43,44^, our study identified the most structured activity in the left anterior hippocampus.

Our findings open up exciting future research in the realms of real-world spatial navigation, episodic memory, and imaginable behaviors. The non-continuous nature of human theta oscillations reported in our study presents an opportunity to delve into the timing and structure of these transient oscillations in various cognitive tasks and behaviors. Subsequent studies could elucidate the neuronal generators of hippocampal theta bouts by simultaneously examining single neurons and local field potentials in freely moving humans^45^, which could shed light on the relationship between distinct functional cell types, theta bouts, and different navigational strategies. Real-world spatial navigation paradigms, in particular, offer a valuable platform for uncovering neural activation patterns relative to task-specific variables. These patterns enable comparisons between navigational and cognitive tasks with similar temporal organization. In studies involving human participants, imaginations can be explicitly instructed and verbally reported, allowing for the investigation of the neural mechanisms underlying abstract, hypothetical, or unprecedented future scenarios, paralleling real-world behaviors.

In summary, our findings highlight the role of transient theta oscillations in partitioning complex navigational routes and episodes into discrete segments, showing a striking similarity in real-world and imagined navigation. This suggests a common neural organization framework underlying both types of navigation independent of environmental constraints. The close resemblance between real-world and imagined navigation aligns with the notion that the neural mechanisms and functional architecture of memory may have evolved from those initially developed for spatial navigation^3,12^. Together with our findings, these parallels underscore the possibility that MTL dynamics contribute to the temporal organization across various cognitive domains, including spatial navigation, episodic memory, and the contemplation of conceivable future scenarios.

## Acknowledgments

We thank all participants for taking part in the study, all members of the Suthana laboratory for discussions, D. Batista for support with the Unity application, and M. Jenkens-Drake for helping with illustrations.

## Funding

National Institutes of Health (NIH) grant U01NS117838 (N.S.)

National Institutes of Health (NIH) grant K99NS126715 (M.St.)

McKnight Foundation, Technological Innovations Award in Neuroscience (N.S.)

Keck Junior Faculty Award (N.S.).

## Author contributions

Conceptualization: M.S., M.St., N.S.

Methodology: M.S., M.V., U.T., N.S.

Investigation: M.S., M.St., M.V., U.T., S.H., C.H., J.P.L., V.R., I.F., D.E., N.S.

Visualization: M.S.

Funding acquisition: N.S.

Project administration: S.H., C.H., J.P.L., V.R., I.F., D.E., N.S.

Supervision: N.S.

Writing – original draft: M.S., N.S.

Writing – review & editing: M.S., M.St., M.V., U.T., S.H., C.H., J.P.L., V.R., I.F., D.E., N.S.

## Competing interests

V.R. is on the Medical Advisory Board for NeuroPace, Inc. All other authors declare no competing interests.

## Data and materials availability

All data and materials used in the analysis supporting the findings of this study will be available before publication.

## Supplementary Materials

Materials and Methods

Figs. S1 to S9

Tables S1 to S2

References (*46*–*54*)

## Supplementary Materials for

### Materials and Methods

#### Participants

Five participants (24-40 years old; three males, two females) who had been chronically implanted with the FDA-approved RNS System (NeuroPace, Inc.) for the treatment of pharmaco-resistant focal epilepsy^27^ volunteered for this study. The electrode placements were determined exclusively by clinical treatment criteria. Individuals with electrodes implanted in the MTL and low rates of epileptic activities were recruited for the study. Prior to participation, all individuals willingly provided informed consent, following a protocol approved by the UCLA Medical Institutional Review Board (IRB).

#### Real-world and imagined navigation tasks

Participants were tasked with traversing two routes (as illustrated in Fig. 1A and B) within an indoor room (measuring 14.6 × 13.5 m^2^), each route featuring four turns. Initially, these route shapes were displayed on a tablet screen. Precise tracking of their walking patterns was achieved through the use of motion capture technology. At the beginning of each recording, participants learned to walk along these patterns without any visible cues about the route’s shapes or the position of the turns. Subsequently, feedback was provided after each walking trial by overlaying their real movement trajectories onto the prescribed ideal routes displayed on the tablet interface, allowing for them to improve their behavior on the next trial. This initial learning phase included the presence of six (three per route) paper cut-out objects positioned along the route. An auditory cue was introduced after random delays during the third segment of each left or right route (following the second turn). Participants were then instructed to determine their relative position to the nearest object by pressing the appropriate left/middle/right button on a wireless handheld mouse, corresponding to the first/second/third object on their route. A second auditory cue signified the accuracy of their response following the button press. This learning phase concluded once participants demonstrated consistent proficiency in navigating these routes. Subsequently, the objects were then removed, requiring participants to rely on memory for walking routes and object locations. Proceeding the learning phase, and during the main experiment, visual feedback was limited, provided only after completing three left and three right walks, serving as regular indicators of their navigation performance.

Real-world navigation trials were interspersed with periods of treadmill walking. These treadmill periods were organized into three distinct blocks over the course of the experiment. Each block consisted of 12 left and 12 right real-world walks, alternating with 24 treadmill walks. Every real-world walk was either preceded or followed by a treadmill walking trial. During the first block of the experiment, participants engaged in treadmill walking devoid of any supplementary tasks. This experimental condition was consistently initiated first, ensuring a baseline of unspecific treadmill walking. Notably, the mention of imagined navigation was omitted to guarantee a naïve experience of sole treadmill walking. In the subsequent two blocks of the experiment (second and third), participants were instructed to mentally navigate their preceding or upcoming walking routes while walking on the treadmill. The utilization of imagined previous and upcoming routes served to investigate the distinction between remembering past spatial trajectories and simulating future ones.

These later two blocks were randomized across participants so that they imagined their previous/upcoming routes during the second/third block, respectively. The imagination periods commenced upon activating the treadmill, and participants signaled the completion of their imagined routes by pressing a button on a wireless handheld mouse. Overall, participants completed a total of 108 walks, including 36 left, 36 right, and 36 imagined walks. The treadmill trials were distributed as follows: 24 imagined left, 24 imagined right, and 24 sole treadmill walks without any imagination involved. These imagination trials were further divided into 24 instances of imagining a previous route and 24 instances of imagining an upcoming route. The exact numbers of trials are listed in Table S1.

#### Unity application

We developed an application using Unity (version 2021.19f1) to control the experimental paradigm, initiate and stop motion tracking recordings, and capture participants’ behavioral responses. In order to facilitate seamless communication, we established a two-way interaction between the Unity application and the motion tracking software (as detailed below). This enabled the Unity application to initiate motion capture recordings and concurrently receive real-time motion data. Motion tracking data was integrated into Unity, enabling the logging of participants’ walking trajectories and positions. This data served as the foundation for visualizing both the actual and ideal walking trajectories, offering participants valuable feedback. For the self-location sub-task, auditory cues were generated based on participants’ positions. Using a wireless handheld mouse, participants’ responses were collected and transmitted, subsequently being recorded and stored within the same Unity application for analysis.

#### Motion capture

To capture participants’ movements, we used the OptiTrack system (Natural Point, Inc.). This setup involved thirty high-resolution infrared cameras strategically positioned on the walls. These cameras recorded the entire laboratory space at a rate of 120 Hz, ensuring precise tracking with sub-millimeter precision. Positions and orientations of the hips, legs, and feet were tracked using a lower-body skeleton fitted to the Helen Heyes marker set. Additionally, head positions and orientation were captured using a rigid body. The gathered motion capture data was analyzed and exported using Motive 3.0 software. Motion capture signals were smoothed with a Gaussian filter using a window size of 0.2 seconds for noise suppression. All walking trajectories are shown in Fig. S1. Movement speed was calculated as positional change in meters per second. Head and hip rotation were quantified as absolute values of angular velocities derived from the head and hip angular orientations (Yaw).

#### iEEG recording

The responsive neurostimulation (RNS) System (NeuroPace, Inc.), approved by the US Food and Drug Administration (FDA), was specifically designed to treat pharmaco-resistant focal epilepsy. This system is engineered to detect epileptic activity and respond in a closed-loop manner by delivering precise electrical stimulation, effectively mitigating the risk of seizures. Notably, the RNS System is implanted chronically, affording the ability to record iEEG activity while individuals move freely (46). In our study, every participant had two depth electrode leads, each comprising four electrode contacts spaced at intervals of 10 mm. We selected specific contacts in each individual based on their location in the MTL to record from up to four bipolar channels in each participant. Data acquisition occurred at a sampling rate of 250 Hz, using the RNS-320 model configured to operate within the broadest available bandwidth, spanning 1-90 Hz.

#### iEEG data analyses

iEEG data was synchronized with motion capture and eye tracking using network time protocol (NTP)-generated markers on all devices^46^. All signals were aligned accordingly and resampled to a frequency of 250 Hz. Time-frequency (TF) analyses were conducted through the utilization of continuous wavelet transform and analytic Morse wavelets, characterized by a symmetry parameter of three and time-bandwidth product set to 60. For frequency bin specification, we employed ten voices per octave within the range of 1 to 90 Hz. Time-varying amplitudes were computed for each frequency. Given the slightly varying durations of real and imagined walking, we employed dynamic time warping to align real walking trials to each other and identify the turns on each walk. This information was then used to linearly adjust motion tracking and time-varying frequency amplitudes, aligning them with the mean timing of all trials. Subsequently, we averaged all trials for left and right walks. The wavelet power was graphically represented in Fig. 2B and utilized to determine frequency spectra in the Fig. S2A through temporal average. Throughout other analyses, we normalized time-frequency amplitudes via z-scoring to account for disparate signal strengths across different recording channels. The theta frequency range was identified for each individual by locating the spectral peak in the 3 to 12 Hz range. The neighboring lower and higher spectral minima determined the exact theta range centered around theta peaks. Individual theta ranges are listed in Table S1.

Time-varying theta amplitudes were rendered on the motion trajectories to directly illustrate their relations to the positions in each walking route. To evaluate if the theta dynamics are temporally structured across trials, we computed the data’s temporal consistency by correlating mean signals from randomly splitting trials in each condition in half. The average values of 1000 random splits for each channel were then statistically compared between conditions (Fig. 3D, Fig. S5A, B). We correlated the temporal consistency values between real-world and imagined walks to investigate whether temporally structured theta dynamics appear at spatially similar sited in both conditions (Fig. 3E).

Individual subject data were aligned the same manner as single trials, using dynamic time warping to match movement trajectories, followed by linear time warping for the time-frequency amplitudes. To present theta dynamics at the group level, the average time and movement trajectories were employed. To analyze the likeness of theta dynamics within each segment of a walking route, we abstracted the navigational task structure with a sinusoidal pattern that peaks at each turn. This approach allowed each segment to possess the same pattern but scaled by its length or relative time. The phase of this sinusoidal pattern corresponded directly to the relative position within a segment. Subsequently, after accounting for segment length discrepancies, we computed the cross-correlation between theta dynamics and the task structure. The central peak of the resulting cross-correlogram served as a metric of the correlation between theta dynamics and the task structure, effectively depicting the geometry and resultant timing across each walking route. Additionally, any side peaks in the cross-correlogram indicated similarities between theta dynamics and shifted task structures, denoting recurring patterns in the data.

By aligning all segments, we were able to calculate the average theta modulation pattern contingent on the relative positions within each segment across all recording channels. Theta dynamics at different electrodes exhibited peaks at marginally distinct relative positions. A modulation akin to a cosine wave would align precisely with the task structure, while a modulation similar to a –sine wave would be orthogonal to it, as depicted in Fig. 4C. The synergy of these two orthogonal components makes them apt for encoding relative position as a circular variable. Leveraging Euler’s formula, the relative position can be mathematically derived as the angle formed between these two orthogonal components.

We established a straightforward linear regression model to examine the feasibility of reconstructing the relative position from theta dynamics. This model encodes the relative segment position as a circular variable, generated through the weighted summation of the theta dynamics of all 18 channels within the MTL. The task structure was used as a response variable to estimate the cosine and –sine modulated components. To prevent overfitting, we used cross-validation (10 folds repeated 10 times) to determine the regression weights. Subsequently, we tested whether the positions estimated from unseen trials aligned with the actual motion capture data. The outcome of this assessment was the computation of estimation errors, which represented the differences between actual and estimated positions.

During real-world navigation trials, we were able to use motion capture to account for minor variations in timing and movement trajectories between trials. However, for the imagined navigation trials, this information was naturally absent. To address this, we instructed participants to mentally navigate the routes in the same manner they did during real-world walking trials. The treadmill played a pivotal role in these trials and conditions, preserving gait patterns without actual progression through space. Rather than relying on motion capture, we recorded the durations between the start and completion of imaginations, which were defined by the onset of the treadmill and the button press reported by participants. During movement, position and time are linearly related through velocity. However, during mental navigation, the imagined velocity of the participant is not directly observable. Imagination trials were aligned via linear time warping for each participant and subsequently averaged across trials. To ensure consistency across the group, participants’ data were adjusted to match the grand average imagination time, yielding a uniform duration. Given the potential variance in imagined velocity profiles across participants and conditions, our aim was to synchronize theta dynamics during imaginations with the navigational task structure of the imagined routes (as depicted in Fig. 4A). Although the exact timing of imagination remained unknown, we capitalized on the structural framework of each imagined route, characterized by five segments connected through four turns.

We utilized the position estimation model developed from our real-world walking data to derive an initial estimate of imagined positions. Subsequently, we used dynamic time warping to align these estimated relative positions with the task structure, allowing for a maximal time shift of ±2 seconds. This alignment procedure was applied for each participant while maintaining the relative timing between recording channels within each participant. In order to prevent overfitting, we employed cross-validation involving 10 folds repeated 10 times to learn the temporal relationship from sub-partitions of the data. This trained alignment was then assessed for its applicability to previously unseen imagination trials, serving as a test of the alignment algorithm’s generalizability. The outcomes of these generalizations are presented throughout the manuscript. It’s worth noting that this alignment procedure remains valid only if the theta dynamics from different imagination trials exhibit similarities. The same procedures for position estimation and time alignment were applied to the sole treadmill walking data, serving as control analyses. By ensuring that results from all conditions were temporally aligned, we were able to calculate position reconstruction errors across all conditions using the actual positions recorded during real-world navigation trials (Fig. 4E). Because the relative position was modeled as a circular variable, we show reconstruction errors as polar histograms of angles between actual and estimated positions.

We detected transient theta oscillations lasting at least two cycles and exceeding the 95% confidence interval of frequency-specific power using the eBOSC toolbox ^47,48^. From these detections, we computed the prevalence for each condition and frequency bin (Fig. S2A) as the percentage of times when a given oscillation was detected. Furthermore, we quantified the percentage of trials in which transient theta oscillations were detected for every time point along a navigational route, resulting in the theta rate (Fig. S2B). To explore the potential influence of eye movements on MTL recordings, we computed average TF activities aligned to the onset of eye saccades and juxtaposed these plots with the ones aligned to turns (Fig. S8). In order to match the number of turns and saccades, four saccadic events were randomly selected in each trial. The temporal relation of behavioral variables and theta dynamics was quantified by calculating cross-correlations of each channel’s theta dynamics and behavioral variables (Fig. S9A, C), averaged across trials.

#### Electrode localization

Post-operative computed tomography (CT) scans were utilized to locate the leads and contacts of the electrodes. To achieve precise anatomical localization for each recording contact, co-registration was performed by aligning the CT scans with pre-operative magnetic resonance images (T1– and/or T2-weighted scans), following a previously established procedure ^49^. Recording sites within the MTL included the hippocampus, amygdala, parahippocampal cortex, and perirhinal cortex (Table S1). Recording contacts situated outside the MTL were excluded from further analyses.

#### Detection of epileptic events

Epileptic events, including interictal epileptiform discharges (IED), were identified using established methods ^50^ and tailored for RNS data (24, 25). In brief, two distinct thresholds were computed, and samples that surpassed either threshold were designated as IED periods and subsequently excluded from further analyses. For this analysis, we computed the envelope of broadband iEEG (1-90 Hz) and filtered (15-80 Hz) signals. The thresholds for IED identification were determined as five times the median values of these signals. To capture both the up and down-ramping epileptic activities before and after IED periods, we extended the identified periods by 256 ms (64 samples) in both directions. In cases where an IED period encompassed over 50% of a trial’s duration, the entire trial was omitted from the analysis. In total, 3.8 ± 3.0% (median ± SD) of the data were identified as IED periods. This relatively low percentage is attributed to our deliberate selection of participants with a limited number of epileptic events per day.

#### Eye tracking

Eye movements were tracked using the Pupil Labs Invisible glasses (Pupil Labs GmbH). The eye data was captured at a frequency of 200 Hz with a resolution of 192 x 192 pixels and synchronized with a scene camera, which sampled the surroundings at 30 Hz, offering a resolution of 1088 x 1080 pixels. To merge these two data streams into coherent gaze position data, we used the Pupil Cloud software ^51^. Saccadic eye movements were identified using the ClusterFix toolbox ^52^. To assess gaze densities, we computed 2D histograms, subsequently smoothed using a Gaussian kernel featuring a standard deviation of 1 pixel. These gaze densities were normalized relative to the expected value of a uniform distribution. The average gaze point in each participant’s visual view was identified to align and center individual gaze densities across participants and conditions. Differences in gaze densities were computed by subtracting gaze densities during left and right real-world and imagined walks (Fig. S7).

#### Statistical comparisons

To quantify the temporal consistency of theta dynamics, we employed a methodology involving the correlation of mean signals derived from randomly dividing the data into two halves. Specifically, trial numbers were randomly grouped into two data sets, the average signals from each group computed, and then linearly correlated using the Pearson correlation coefficient. This procedure was repeated 1000 times to yield a robust measure of how theta dynamics from one randomly selected half of the trials were associated with the other half. The experimental conditions were compared using multi-level block permutation tests^53,54^ while restricting exchangeability blocks to each participant (Fig. 3D, Fig. S5A, B). The temporal consistency values for each channel were then subjected to correlation to explore the presence of temporally structured theta dynamics at spatially similar locations across different conditions. The statistical significance of this correlation was evaluated using linear mixed effect models using the participant numbers as a grouping variable (Fig. 3E). These tests were applied to account for the grouped nature of our recordings consisting of multiple electrodes grouped per participant.

Nonparametric permutation tests were employed to determine the significance of theta amplitudes dependent on positions along the routes. Multiple comparisons were corrected using cluster-level statistics ^53^. First, the mean theta dynamics across channels were computed. Second, a permutation distribution was obtained by sign flipping of randomly selected half of the channels and subsequent averaging, repeated 1000 times. Third, a primary threshold of p = 0.05 was applied to identify clusters of significant activity. Fourth, these cluster sizes were compared against the clusters of the permutation distribution and were only considered significant when surviving a secondary p = 0.05 (cluster) threshold correcting for multiple comparisons (Fig. S4B).

We utilized nonparametric permutation tests to compare the position reconstruction errors against the control condition, which was sole treadmill walking. Specifically, we randomly re-assigned the labels of reconstruction errors for the real-world and control conditions and averaged them 10,000 times, resulting in a permutation distribution. The actual reconstruction errors were then compared to this permutation distribution. The same procedure was analogously applied to the imagined navigation data. The probabilities of reconstructed positions (as depicted in Fig. 4D) were normalized relative to the expected value of a uniform distribution. This scale indicates by which factor the position reconstruction results exceed this chance distribution.

We tested for a significant correlation between theta dynamics of real-world and imagined navigation of trials generalizing the alignment procedure. This correlation was compared to values computed from randomly shifting the real– and imagined navigation data 10,000 times to disrupt their temporal relationship (Fig. S5C). The same analyses were performed to assess the correlation of theta dynamics and the maze structure (Fig. 4D).

Additionally, we computed linear regression models of theta dynamics and behavioral variables on a single trial level for each recording electrode. Subsequently, we tested the effects on the group level and compared the resulting regression coefficients using multi-level block permutation tests while controlling for multiple comparisons ^53,54^.

**Fig. S1.**
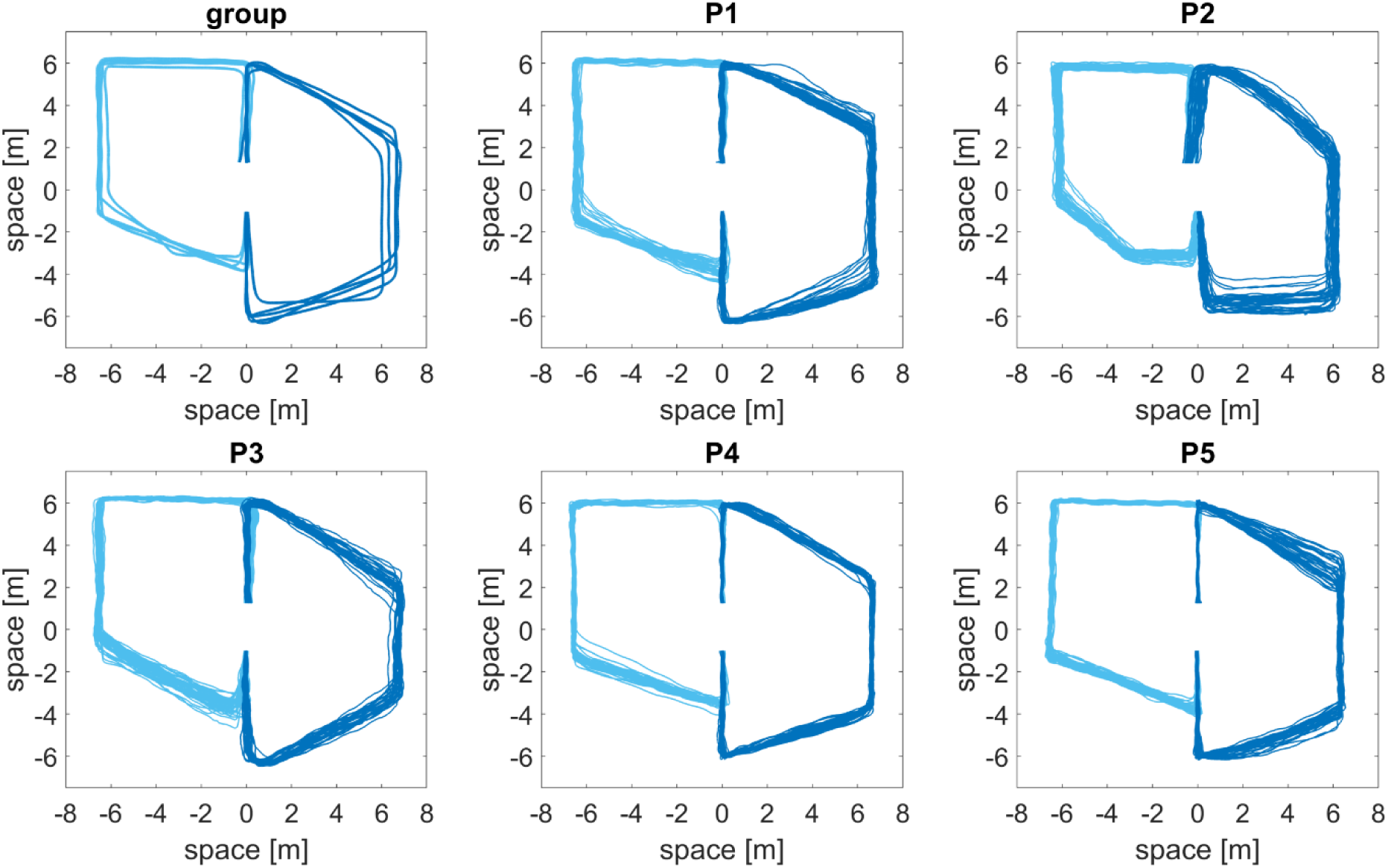
| Motion capture data. Average walking routes (N = 5 participants) of each participant shown in a group plot (top left). Walking trajectories of each trial for every participant (P1 P5) shown in light/dark blue for the left/right route, respectively.

**Fig. S2.**
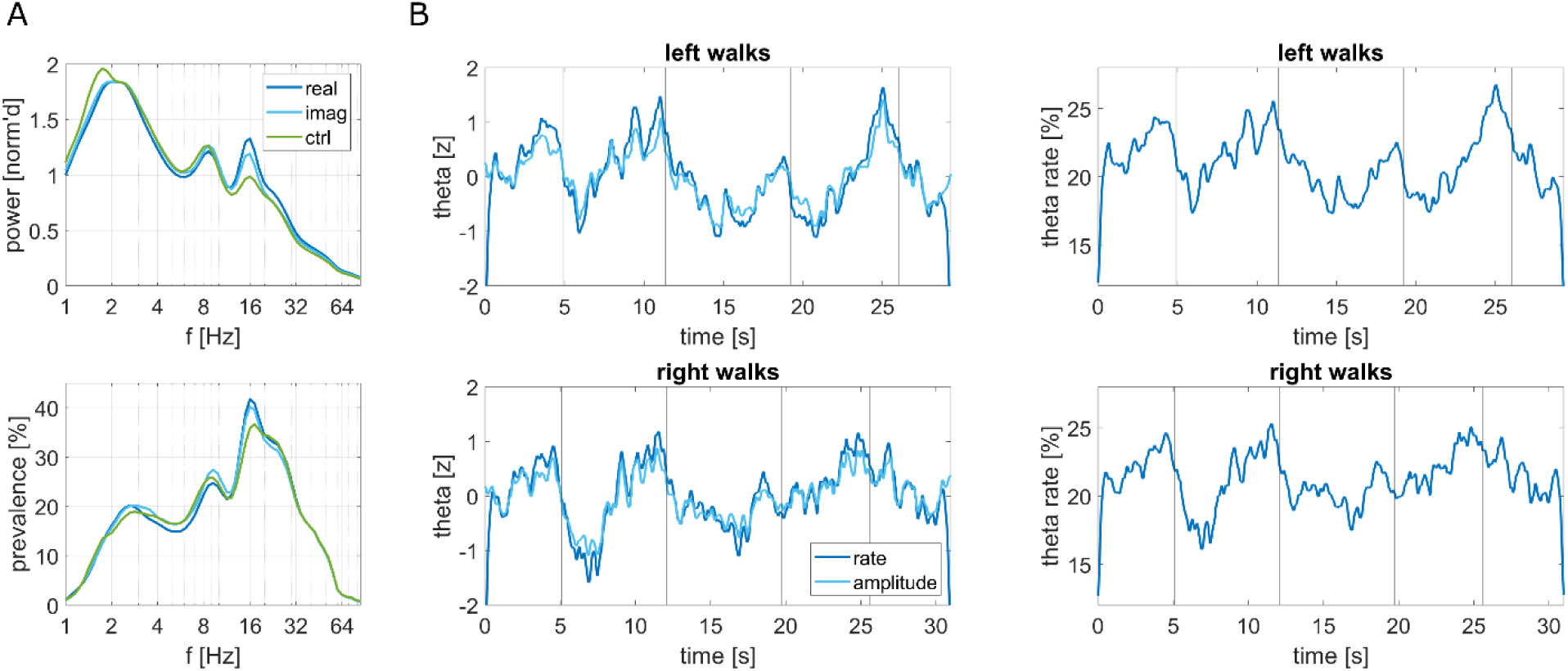
| Power spectra and theta dynamics. (**A**), Normalized power spectra averaged across participants for real-world, imagined navigation and the control condition (top). Prevalence of oscillatory bouts (bottom). (**B**), Theta amplitude dynamics and time-resolved theta rates z-scored for comparison (left), and theta rates shown as detection percentages across trials (right) for left (top) and right (bottom) walks.

**Fig. S3.**
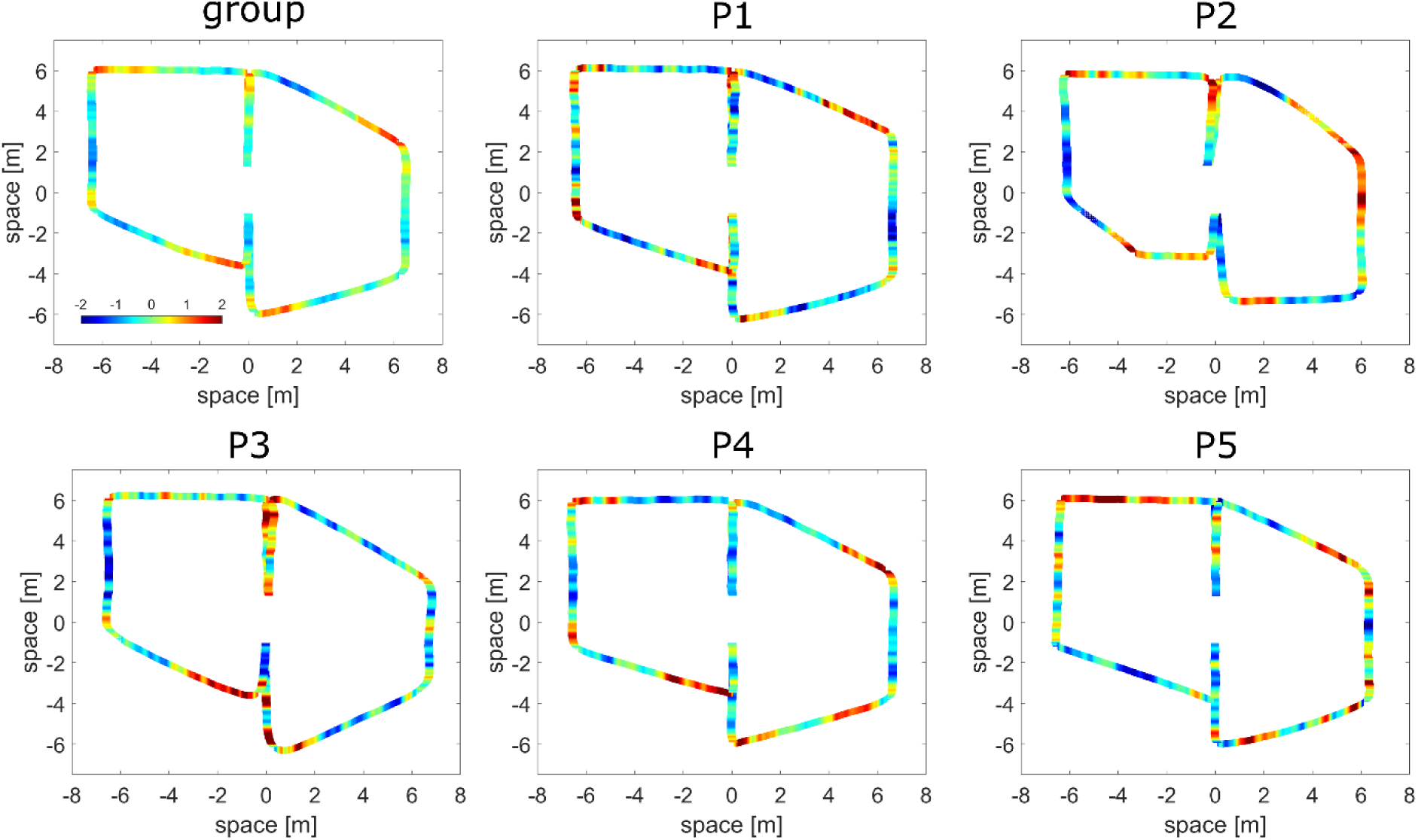
| Individual theta activity patterns. Mean group and individual participant (P1-5) theta activity patterns within the left anterior hippocampus (top left panel) superimposed onto the mean behavioral motion trajectories, illustrating an increase in theta power as each of the participants approaches upcoming turns.

**Fig. S4.**
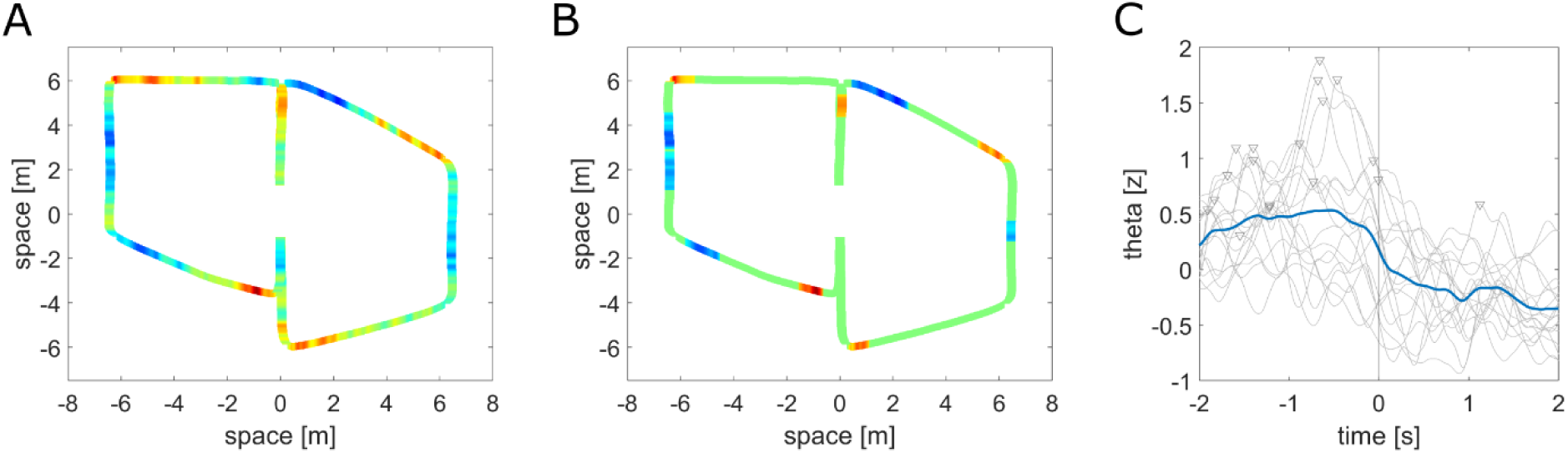
| Theta activity and timing of effect. (**A**), Superimposed averaged theta activity across all channels on the mean motion trajectories. (**B**), Statistically significant theta clusters (p < 0.05, determined through a cluster-based permutation test). (**C**), Average theta activity aligned with turns depicted in blue, while individual electrode data is presented in grey. Triangles indicate that peak theta activity precedes turns in 17 out of 18 channels.

**Fig. S5.**
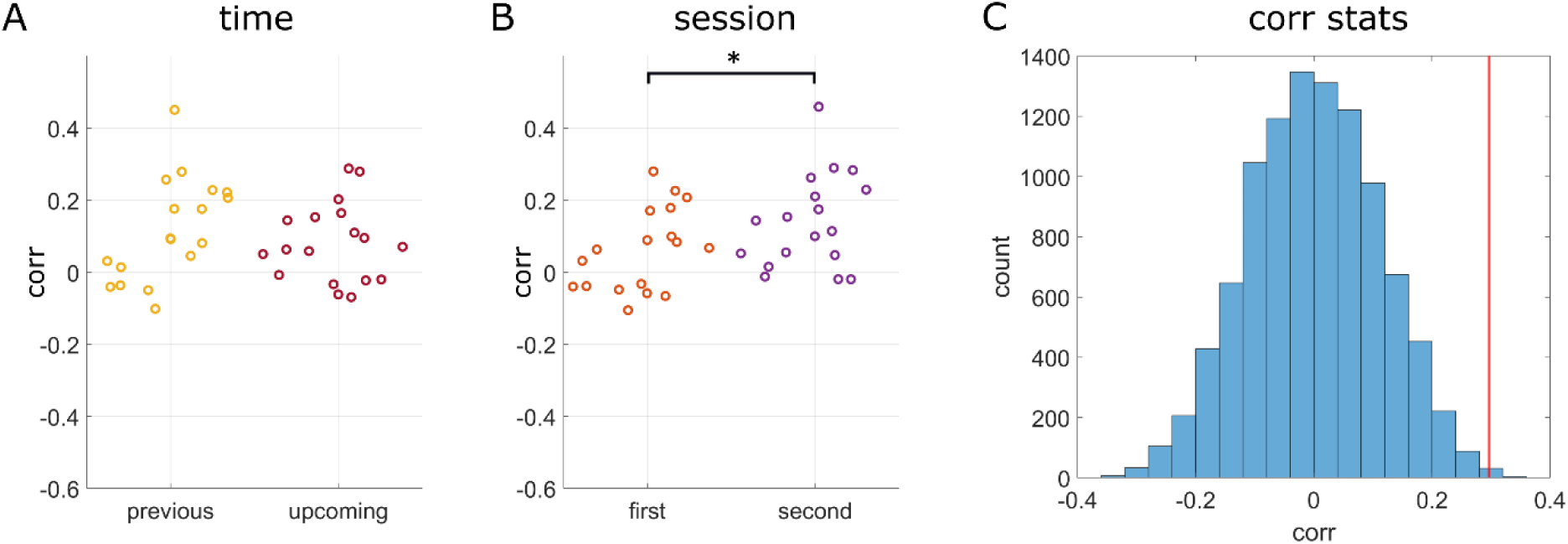
| Theta dynamics across conditions. (**A**), Temporal consistency, measured using a correlation coefficient (corr), during imagination of previous (past) and upcoming (future) routes. (**B**), Temporal consistency observed within the first and second imagination sessions. (**C**), Permutation distribution and actual correlation value between theta dynamics of real– and imagined navigation (red line).

**Fig. S6.**
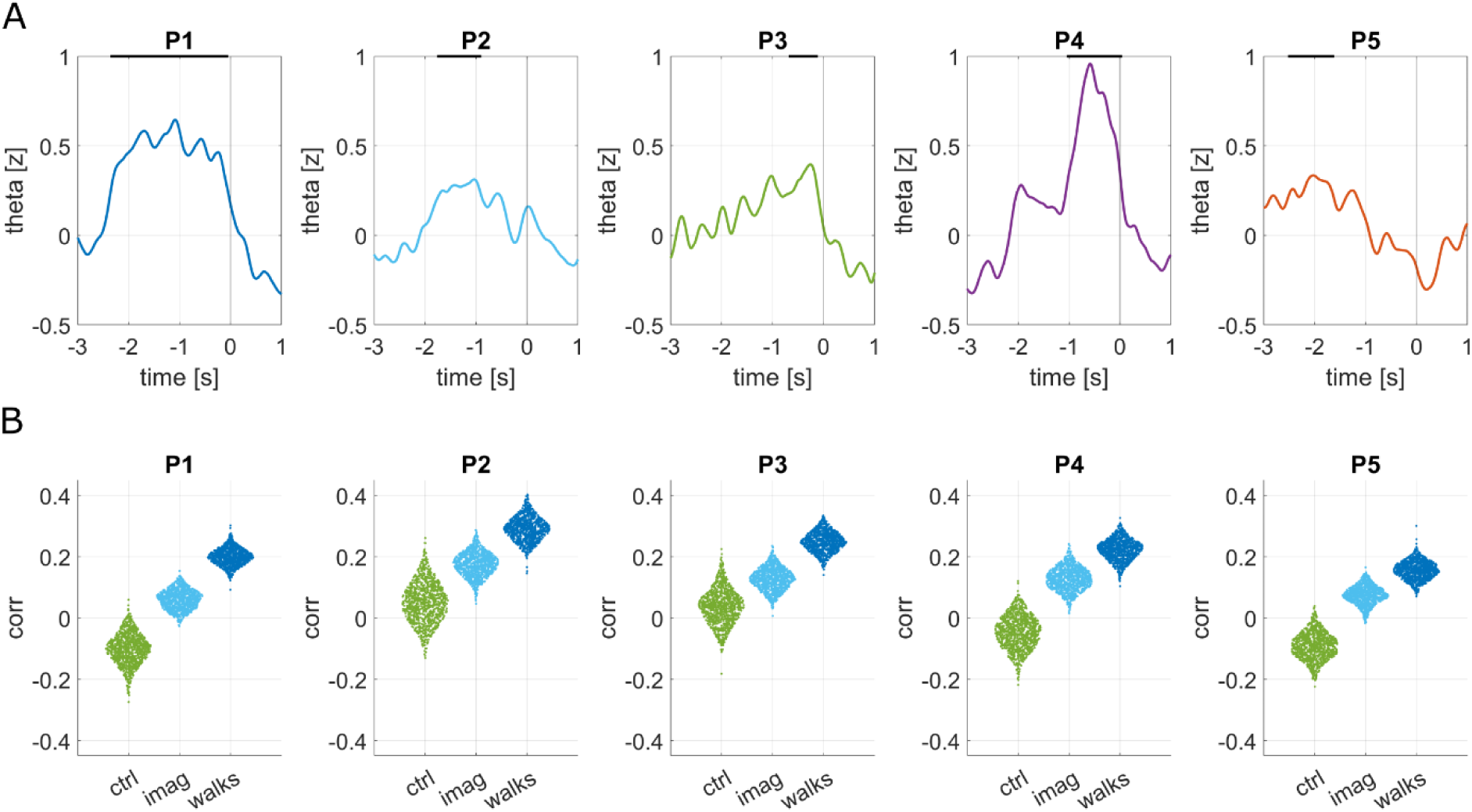
| Individual subject analyses. (**A**), Mean theta activity across channels for each participant (P1-P5) aligned to turns (t = 0). Black bars on top of the plots indicate significant periods. (**B**), Temporal consistency assessed for each participant. Individual circles represent data from correlations computed by randomly half-split data a thousand times. Differences between conditions are significant (p < 0.001) in every subject.

**Fig. S7.**
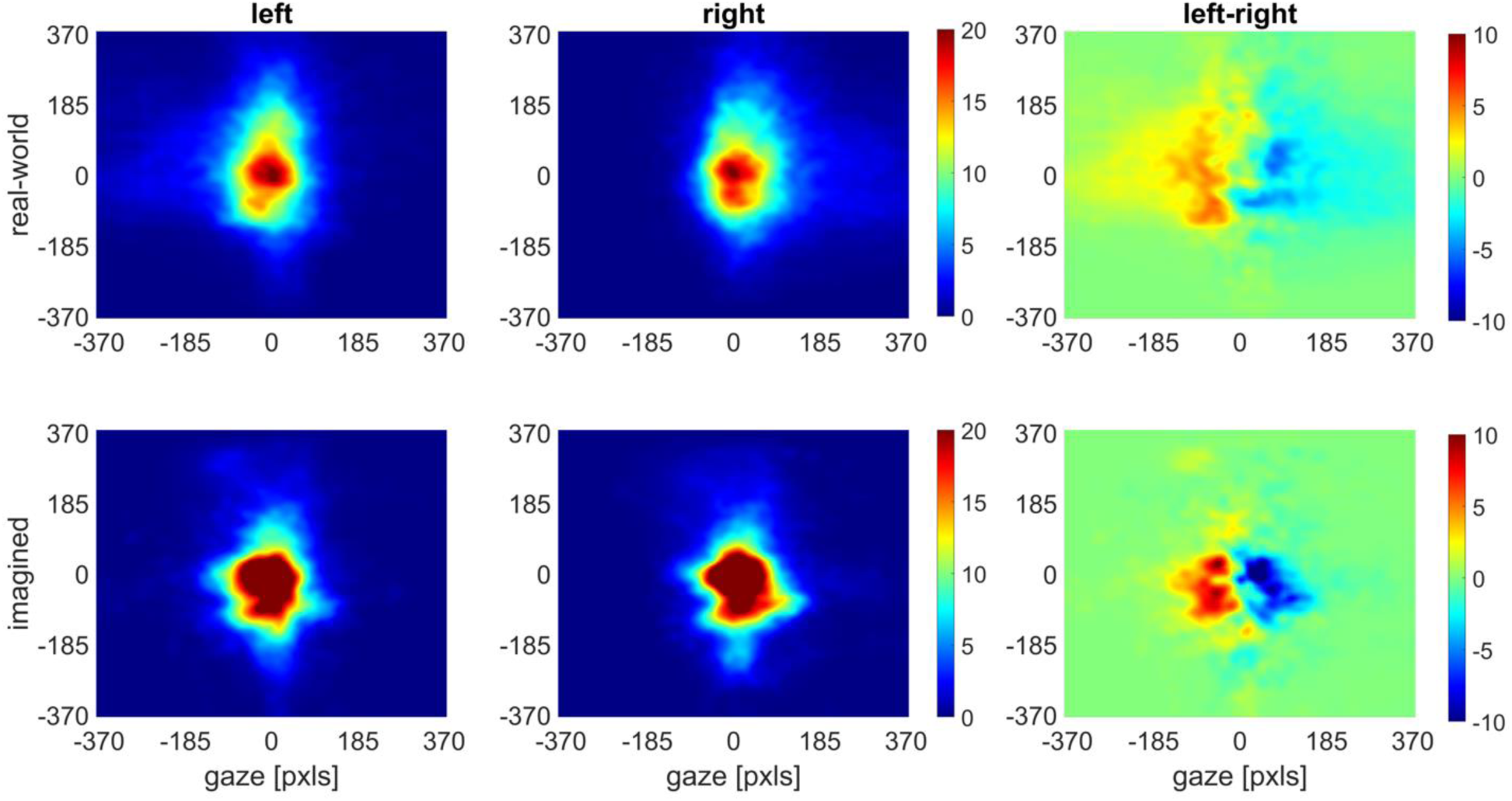
| Gaze densities. Shown are the 2D histograms, centered at participants’ natural fixation point within their field of view. During real-world navigation on the left or right route, participants tended to direct their gaze more towards the respective left or right directions. Similarly, during imagined navigation, participants exhibited a slight bias in their gaze towards the left or right visual field, depending on whether they were simulating a left or right route. This pattern of gaze behavior confirmed their active engagement in the task.

**Fig. S8.**
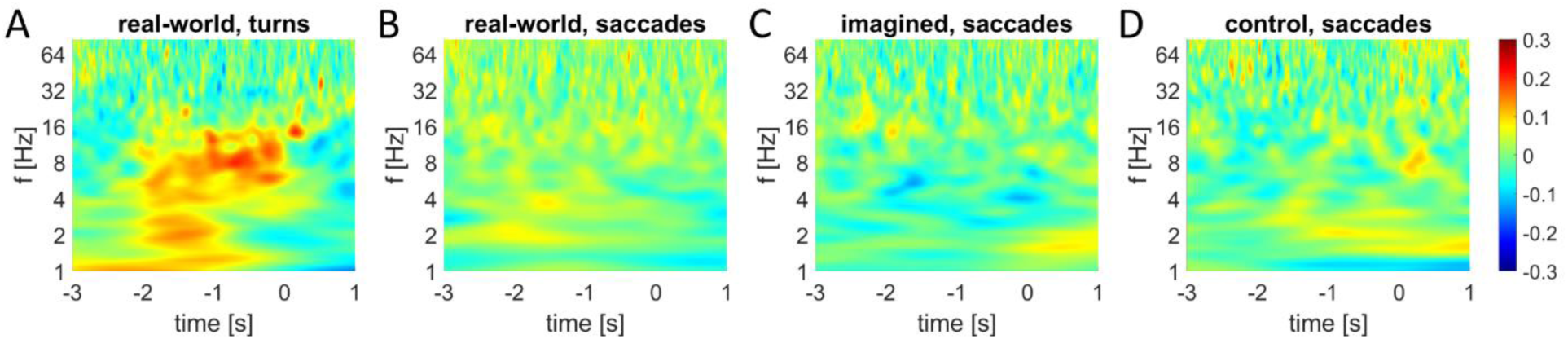
| MTL oscillatory activity aligned to turns and saccades. Time-frequency activity aligned to turns (**A**) and saccades (**B**) during real-world navigation. Time-frequency activity aligned to eye movements (saccades) during imagined navigation (**C**) and the control navigation condition of treadmill walking without imagination (**D**). Note that the well-pronounced theta activities precede physical turns but not saccades.

**Fig. S9.**
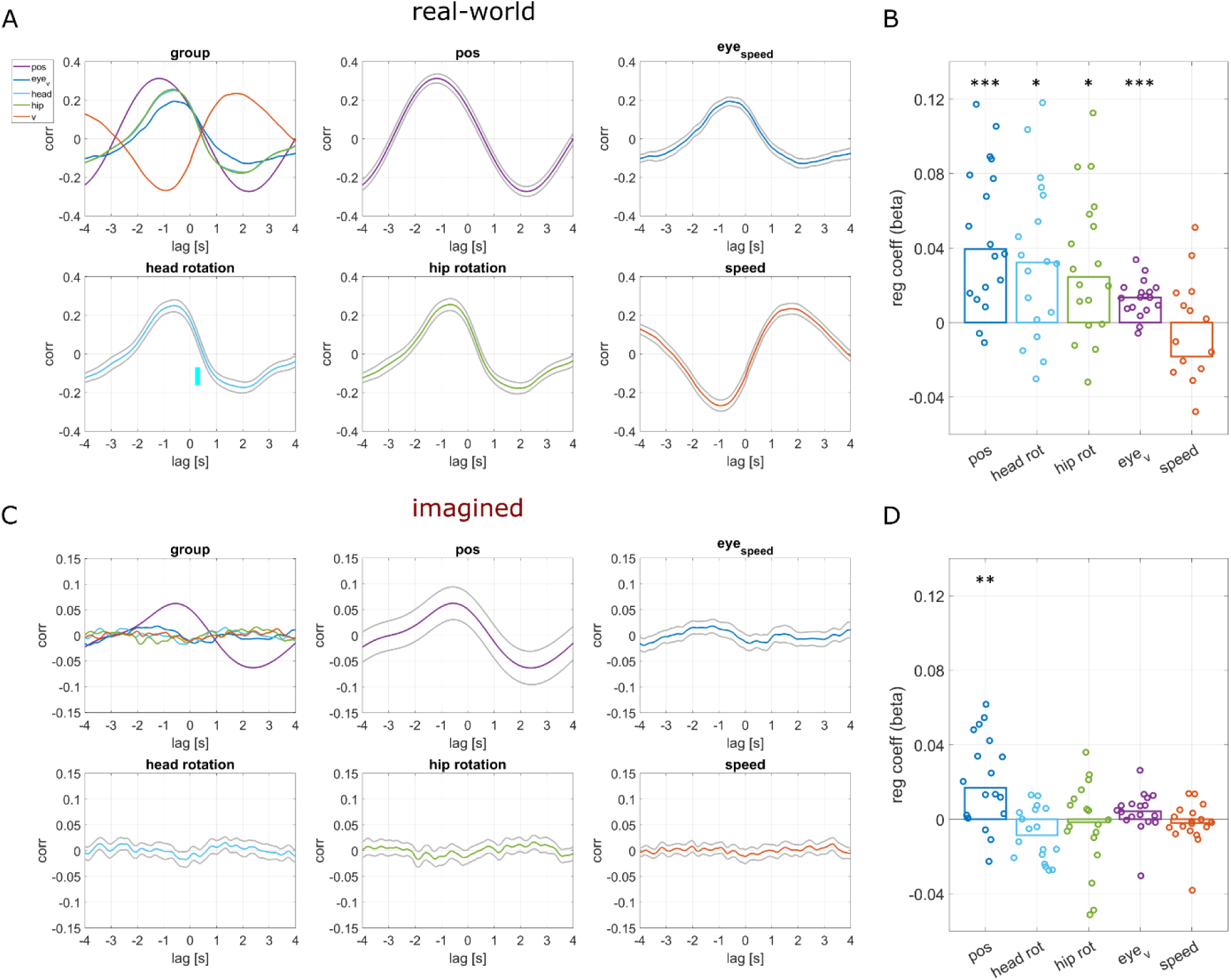
| Relation of theta dynamics to behavioral variables. (**A**), Cross-correlation of theta dynamics and behavioral variables during real-world navigation. Colored lines illustrate the group average, while grey lines represent standard errors across the folds of cross-validation for each behavioral variable (position [pos], eye speed, head rotation, hip rotation, and movement speed [v]). Negative lags indicate that theta dynamics preceded the behavioral signals. An overview of temporal relationships for each behavioral variable is summarized in the top left panel, showing that theta activities peaked at approximately 1 second before physical turns, followed by head rotation, eye saccades, speed reductions and hip rotation. The positive lagged correlation between theta and movement speed can be attributed to the speed decrease before turns, followed by a reduction in theta activity. (**B**), Regression coefficients of ongoing (single trial) theta dynamics during real-world navigation with labeled behavioral variables. (**C**) Cross-correlation of theta dynamics and behavioral variables during imagined navigation analogous to (**A**). Theta dynamics are compared to behavioral variables after alignment to enable comparisons to position within the maze segments. (**D**), Regression coefficients of ongoing (single trial) theta dynamics during imagined navigation as in (**B**).

**Table S1.**
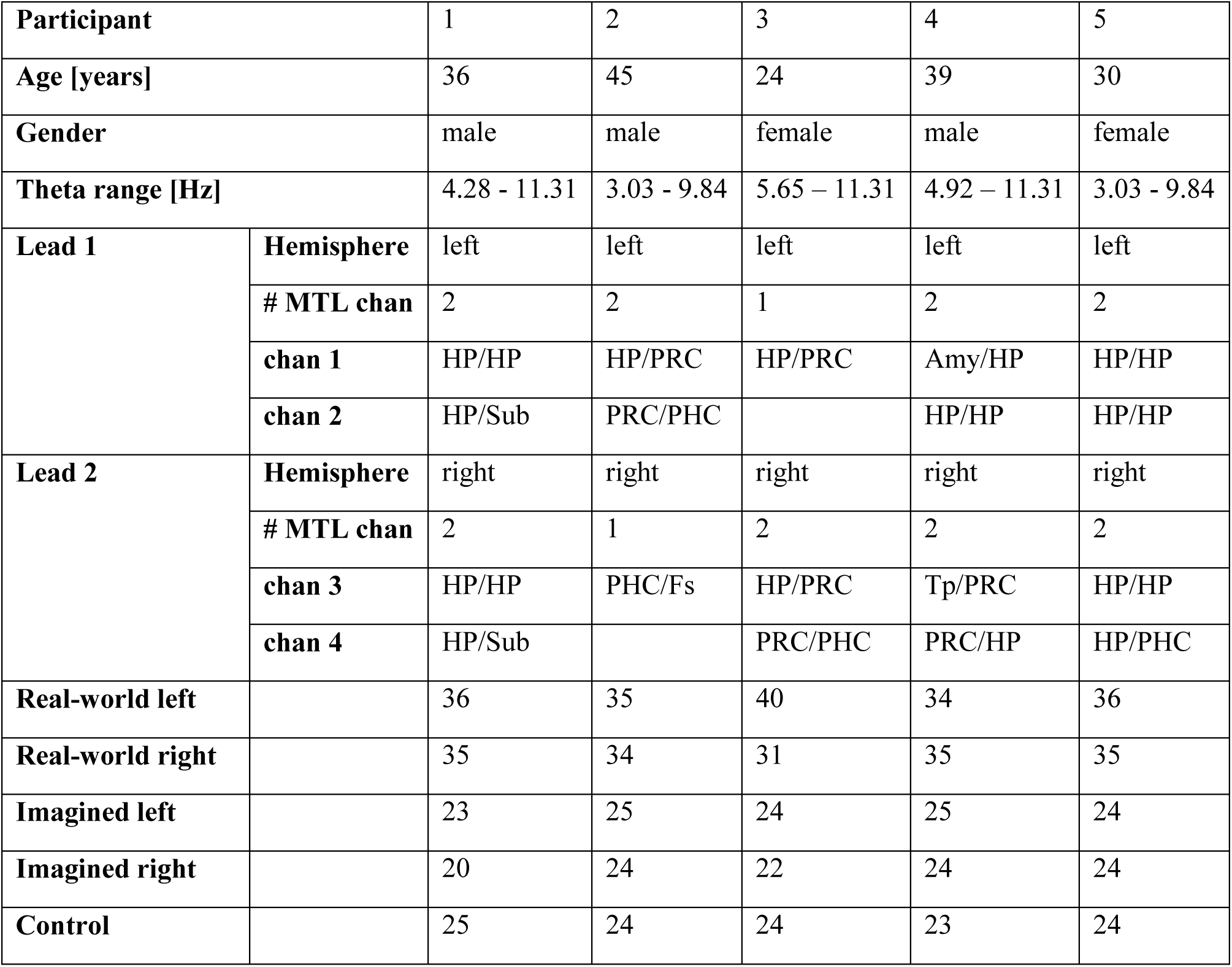
| Participants and experimental information. Participants demographics, theta frequency ranges, electrode implantation sites, number of trials completed for real-world left and right walks, imagined left and right walks, and treadmill control walks. Theta range was determined by spectral peaks in the 3-12 Hz range and neighboring lower and higher spectral minima. Localization of each of the two contacts that form a bipolar channel (chan 1 and 2) in various medial temporal lobe (MTL) regions, including the hippocampus (HP), perirhinal cortex (PRC), parahippocampal cortex (PHC), subiculum (Sub), Fusiform gyrus (Fs), temporal pole (Tp), and amygdala (Amy).

**Table S2.** | Statistics of theta dynamics and behavioral relations. Statistics (p-values) testing whether theta dynamics precede behavioral variables (Lag) during real-world navigation are listed in the top row. Relations between theta dynamics and behavioral variables were assessed on a single-trial level and tested for differences from zero on a group level. The results of these tests (p-values) are listed for real-world (middle row, corresponding to Fig. S9B) and imagined navigation (bottom row, corresponding to Fig. S9D). Multi-level block permutation tests considered data grouping and corrected for multiple comparisons.

|  | **pos** | **head rot** | **hip rot** | **eye_v** | **speed** |
| --- | --- | --- | --- | --- | --- |
| **Lag** | **0.0001** | **0.0001** | **0.0001** | **0.0021** | **0.0001** |
| **Walks** | **0.0001** | **0.0123** | **0.0174** | **0.0001** | 0.1503 |
| **Trdm** | **0.0063** | 0.0508 | 0.9816 | 0.5676 | 0.9194 |

